# NK cell allorecognition shapes reprogramming of neutrophils infiltrating heart allografts

**DOI:** 10.1101/2025.09.28.679079

**Authors:** Erik H. Koritzinsky, Karen S. Keslar, Robert L. Fairchild, Jayati Basu

## Abstract

**Background:** Recent developments in neutrophil biology have demonstrated that neutrophils are phenotypically and functionally heterogeneous. Tissue microenvironments dictate changes in neutrophil cell states in infection and cancer, but little is known about how alloimmune responses or solid organ transplantation influence neutrophil heterogeneity and plasticity.

**Methods:** Here, we used the murine heterotopic heart transplant model in conjunction with high dimensional flow cytometry and transcriptome analysis to interrogate how the alloimmune response and microenvironment of a transplanted organ influence neutrophil subset differentiation and plasticity. Complete MHC mismatched A/J (H2^a^) or syngeneic B6 (H2^b^) hearts were transplanted to C57BL/6 or B6 background genetically modified recipients.

**Results:** We uncovered striking differences between neutrophils infiltrating complete MHC mismatched allografts and syngeneic isografts. Bone marrow neutrophil development was highly skewed towards an immature, interferon stimulated gene (ISG)^+^ subset (marked by IFIT1 expression) early after transplant in both allo- and iso-graft recipients. ISG^+^ neutrophils were also the dominant population in the peripheral blood of both recipient groups. In contrast, neutrophils maintained the ISG^+^ phenotype after infiltrating an allograft but appeared to turn off this program upon infiltrating an isograft. The heart graft microenvironment imposed additional reprogramming independent of donor-recipient mismatch, as neutrophils from both allo- and iso-grafts were skewed towards a mature, aged and proangiogenic dcTRAIL-R1^+^ phenotype. Interestingly, while the existing literature indicates that IFIT1-expressing ISG^+^ neutrophils and proangiogenic dcTRAIL-R1^+^ neutrophils are distinct subsets, we identified a novel IFIT1^+^ dcTRAIL-R1^+^ hybrid population that is highly enriched in allografts. Mechanistically, NK cell-mediated innate allorecognition drives this early intra-allograft specific neutrophil phenotypic programing.

**Conclusions:** These findings provide novel insights into the innate immune allorecognition-mediated regulation of the plasticity of recently described key neutrophil subsets and will enable specific targeting to neutralize detrimental neutrophil subsets and enhance solid organ transplant outcomes.

## Introduction

Solid organ transplantation remains the optimal treatment strategy for end stage organ diseases. Organs harvested for transplant are subjected to periods of cold ischemic storage (CIS) during transport and warm ischemia during surgery. These factors are key determinants of the extent of ischemia-reperfusion injury (IRI) that occurs after revascularization in the recipient. Clinically, the magnitude of IRI negatively impacts short- and long-term graft outcomes.^1–3^ Increased IRI mediates early infiltration and activation of innate immune cells that promote memory CD8^+^ T cell activation to cause acute graft injury and CTLA-4Ig costimulatory blockade resistant acute graft rejection.^4–6^ Therefore, adjunct therapies targeting early infiltrating innate immune cells may enhance the success of costimulatory blockade-based immunosuppressive regimens.

Recent developments in the field of neutrophil biology have revealed significant phenotypic and functional neutrophil heterogeneity in homeostasis, malignancy and infection.^7–11^ Importantly, this heterogeneity is not fixed. Neutrophils are highly plastic and their cell states can be differentially modulated upon entering the microenvironment of different organs.^12,13^ Evidence from single-cell omics data suggests that circulating neutrophils are phenotypically and functionally diverse. One key study identified a pro-inflammatory subset (G5b) that rapidly expands during infection and is marked by the expression of interferon stimulated genes (ISGs), including IFIT1.^10^ The G5b/ISG phenotype is generally programmed in the bone marrow (BM). Beyond the BM, further neutrophil specialization and reprogramming can occur in the context of different tissue microenvironments.^12,13^ For example, the tumor microenvironment has been shown to reprogram infiltrating neutrophils to express tumor growth promoting proangiogenic capabilities, marked in mice by cell surface decoy TRAIL receptor 1 (dcTRAIL-R1) expression.^11^ Currently, little is known about neutrophil heterogeneity in the response to solid organ transplantation.^14^ Most importantly, whether and how IRI couples with donor-reactive immune responses to influence neutrophil phenotypic programing in a transplanted organ has not been investigated.

Neutrophils are the first leukocyte responders to IRI. Data from mouse heterotopic heart transplant models show that neutrophils infiltrate solid organ grafts within minutes of reperfusion^15^, with the influx peaking at 24h post-transplant.^16,17^ We have shown that targeting early neutrophil infiltration in this model via anti-CXCR2 antisera, anti-CXCL1+CXCL2 antisera, or anti-GR1 depleting antibody treatment reduces early allograft inflammation and T cell infiltration, and can prolong allograft survival when coupled with costimulatory blockade.^16^ Clinically, peri-transplant myeloablation or global inhibition of neutrophil function is not a viable option, as these would further increase the already high risk of early post-surgery infection, a comorbidity that is associated with poorer short- and long-term transplant outcomes.^18–20^ Despite our pre-clinical heart transplant data, subsequent work using a mouse myocardial infarction model suggests that neutrophils facilitate effective myocardial repair and enhance post-ischemic recovery of heart function following myocardial infarction.^21^ These seemingly paradoxical findings about the role of neutrophils in responding to cardiac ischemia might be explained by differences in subset specific functions, plasticity and/or developmental programming of neutrophils, which are all poorly understood.

Here, we used the heterotopic heart transplant model in conjunction with high dimensional flow cytometry and gene expression analysis to interrogate how the alloimmune response and microenvironment of a transplanted organ influence neutrophil subset differentiation and plasticity. We show that BM neutrophil development was highly skewed towards an immature, ISG^+^ subset (marked by IFIT1 expression) early after transplant in both allo- and iso-graft recipients. ISG^+^ neutrophils were also the dominant population in the peripheral blood of both groups of recipients. Upon entering allo- or iso-heart grafts, early infiltrating neutrophils also displayed a mature, aged, and pro-angiogenic dcTRAIL-R1 expressing phenotype. Further, we uncovered striking differences between neutrophils infiltrating complete MHC mismatched allografts and syngeneic grafts. Neutrophils maintained the ISG^+^ phenotype (IFIT1^+^) after entering an allograft but appeared to turn off this program upon entering an isograft (IFIT1^-^). Interestingly, while the existing literature indicates that ISG^+^ neutrophils and proangiogenic dcTRAIL-R1^+^ neutrophils are distinct subsets^11^, we identified a novel IFIT1^+^ dcTRAIL-R1^+^ hybrid population that is highly enriched in allografts. Mechanistically, we show that recipient NK cells drive this early allograft specific neutrophil phenotypic programing. These findings provide novel insights into the regulation of recently described key neutrophil subsets that could enable more specific targeting of detrimental neutrophil subsets to enhance solid organ transplant outcomes.

## Materials and Methods

### Mice

C57BL/6J (H-2^b^, Strain 000664), A/J (H-2^a^, Strain 000646) and B6.129S7-*Rag1^tm1Mom^*/J (*Rag1^-/-^*, Strain 002216) mice were purchased from The Jackson Laboratory (Bar Harbor, ME). C57BL/6NTac.Cg-*Rag2^tm1Fwa^ Il2rg^tm1Wjl^* (*Rag2^-/-^IL2Rg^-/-^*, Model No. 4111) mice were purchased from Taconic Biosciences (Germantown, NY). All mice were housed in specific-pathogen free conditions with ad libitum access to food and water in the Cleveland Clinic Lerner Research Institute Biological Resources Unit. All animal studies were approved by the Cleveland Clinic Institutional Animal Care and Use Committee under protocols 2143 and 2174.

### Heterotopic heart transplantation

Murine intra-abdominal heterotopic heart transplantation was performed as adapted from Corry et al.^22^ Allogeneic (complete MHC mismatched) or syngeneic donor hearts were stored in Ringer’s solution (R5310-01, B. Braun Medical Inc., Bethlehem, PA) on ice before implantation in the recipient. The donor graft aorta and pulmonary artery were anastomosed to the recipient abdominal aorta and inferior vena cava respectively. Graft viability was verified after closure by abdominal palpation. All recipients were unsensitized. All heart donors and recipients were 8-12 week old male kept in mice specific-pathogen free caging. Grafts that stopped beating within the first 72 hours post-transplant were considered technical failures and excluded from analysis.

### Processing of tissues to obtain cells for flow cytometry analysis

#### Bone Marrow

Femurs and tibias were harvested, the tips were cut and the bones were flushed with RPMI to collect the marrow cells. The suspensions were then diluted with additional RPMI before filtering through a 40 µM filter. The cell suspensions were spun down and resuspended in 1X RBC Lysis Buffer (420301, Biolegend, San Diego, CA) for 5 minutes before counting and additional processing for downstream use.

#### Blood

Blood was collected and diluted 1:10 in heparinized PBS. Samples were spun down and resuspended in 1X RBC Lysis Buffer (420301, Biolegend, San Diego, CA) for 5 minutes before counting and additional processing for downstream use.

#### Heart grafts

After perfusion with cold Ringer’s solution, harvested heart grafts were weighed, minced, and incubated in RPMI with 1 mg/ml Collagenase D (11088882001, Roche, Mannheim, Germany) and 0.06 mg/ml DNase I (10104159001, Roche, Mannheim, Germany) for 45 minutes at 37°C in a thermal mixer at 750 RPM. The digested tissue was pressed through a 70µM filter with the plunger of a syringe, diluted with additional RPMI, and filtered through a 40 µM filter. The cell suspensions were spun down and resuspended in 1X RBC Lysis Buffer (420301, Biolegend, San Diego, CA) for 5 minutes before counting and additional processing for downstream use.

#### Spleen

Spleens were collected in RPMI and pressed through a 70µM filter with the plunger of a syringe, diluted with additional RPMI, and filtered through a 40 µM filter. The cell suspensions were spun down and resuspended in 1X RBC Lysis Buffer (420301, Biolegend, San Diego, CA) for 5 minutes before counting and additional processing for downstream use.

### Flow cytometry staining and analysis

Cells were incubated with mouse Fc Block (anti-mouse CD16/32 clone 2.4G2, 553141, BD Biosciences, San Jose, CA) before adding conjugated antibodies for surface staining. Antibodies were added to Brilliant Stain Buffer Plus (566385, BD Biosciences, San Jose, CA) to make cocktails before adding to samples. Samples stained with a biotin conjugated primary antibody were washed after with 1X PBS/4% FBS (HyClone Fetal Bovine Serum: SH30070.03, Cytiva, Marlborough, MA) before incubation with a streptavidin conjugated secondary antibody. Cells were then washed with protein-free 1X PBS before incubation with LIVE/DEAD Fixable Aqua stain (L34965, ThermoFisher Scientific, Waltham, MA). Samples were then fixed and permeabilized for 30 minutes on ice using the eBioscience Foxp3/Transcription Factor Staining Buffer Set (00-5523-00, ThermoFisher Scientific, Waltham, MA). For intracellular staining, antibodies were diluted in Foxp3/Transcription Factor Permeabilization Buffer and incubated for 1hr at room temperature. After final washes, data from stained cells was acquired using a BD LSR Fortessa-X20 flow cytometer (BD Biosciences, San Jose, CA). Flow cytometry data was analyzed with FlowJo v10.10 (BD Biosciences, San Jose, CA). MFIs are reported as Median Fluorescence Intensity. All antibodies used for flow cytometry staining are listed in the Major Resources Table in the Supplementary Material.

### Ex vivo stimulation of heart graft cells

Aliquots of cell suspensions from heart grafts were stimulated using eBioscience Cell Stimulation Cocktail plus protein transport inhibitors (00-4975-93, ThermoFisher Scientific, Waltham, MA) in complete RPMI for 4 hours. Cells were then stained with a fixable viability stain and then fixed, permeabilized and stained as described above.

### RNA extraction and NanoString nCounter analysis

Total RNA was extracted from pieces of harvested heart grafts using the RNeasy Fibrous Tissue Mini Kit (74704, QIAGEN, Germantown, MD). One hundred ng of RNA from each sample was hybridized to the nCounter PanCancer Immune Profiling Codeset (NanoString, Bruker Spatial Biology, Seattle, WA) and then processed on a GEN2 analysis system using the high sensitivity protocol and high resolution data capture. Raw data was uploaded to the Rosalind analysis platform (https://app.rosalind.bio) to generate Log2 normalized counts and expression ratios and perform statistical analysis for differentially expressed genes as originally implemented in nSolver Advanced Analysis software. Cutoffs of *p* ≤ 0.05 and a minimum 2-fold change were used to identify genes that were significantly differentially expressed between two groups.

### Enrichr pathway analysis

Pathway analysis on sets of unique or shared differentially expressed genes was performed using Enrichr (https://maayanlab.cloud/Enrichr)^23–25^ based on the Reactome^26^ pathways library. Adjusted *p* values were calculated using the Benjamini-Hochberg method for correction for multiple hypotheses testing.

### Statistical analysis

Statistical analysis was performed using GraphPad Prism 10 (GraphPad Software, Boston, MA). Comparisons between two groups were analyzed by two-tailed Students’ *t*-test and comparisons involving three or more groups were analyzed by one-way ANOVA with Šídák’s multiple comparisons test. A *p* value threshold of <0.05 was chosen as the cutoff for statistical significance. All data are expressed as mean ± SEM. Analysis of Nanostring RNA expression data is described above.

### Data availability

The complete Nanostring dataset from Figure 4 has been deposited in the NCBI Gene Expression Omnibus (GEO XXXXX). All other data are available upon reasonable request.

## Results

### Graft infiltrating neutrophils are distinct from recipient blood and bone marrow neutrophils

Considering the limited understanding of neutrophil diversity during transplant ischemia-reperfusion injury, we performed a detailed analysis of immune cell populations in murine heterotopic heart grafts transplanted to unsensitized recipients on day 1 post-transplant and compared them to populations in matched recipient BM and blood. Comparing complete MHC mismatched A/J (H2^a^) allografts with syngeneic B6 (H2^b^) isografts allowed us to dissect the impact of the alloimmune response from that of transplant IRI. We found that neutrophils represent >50% of heart allo- and iso-graft infiltrating immune cells at 24h post-transplant. However, we observed only a marginal difference in the frequency and no significant difference in the number of infiltrating neutrophils between allo- and iso-grafts (Fig 1A-B). There was also no significant difference in neutrophil frequency or number in the BM or blood between allo- and iso-graft recipients (Fig S1). The next largest populations of graft immune cells were macrophages and NK cells, followed by minimal numbers of T and B cells at this early timepoint (Fig S2).

**Figure 1.**
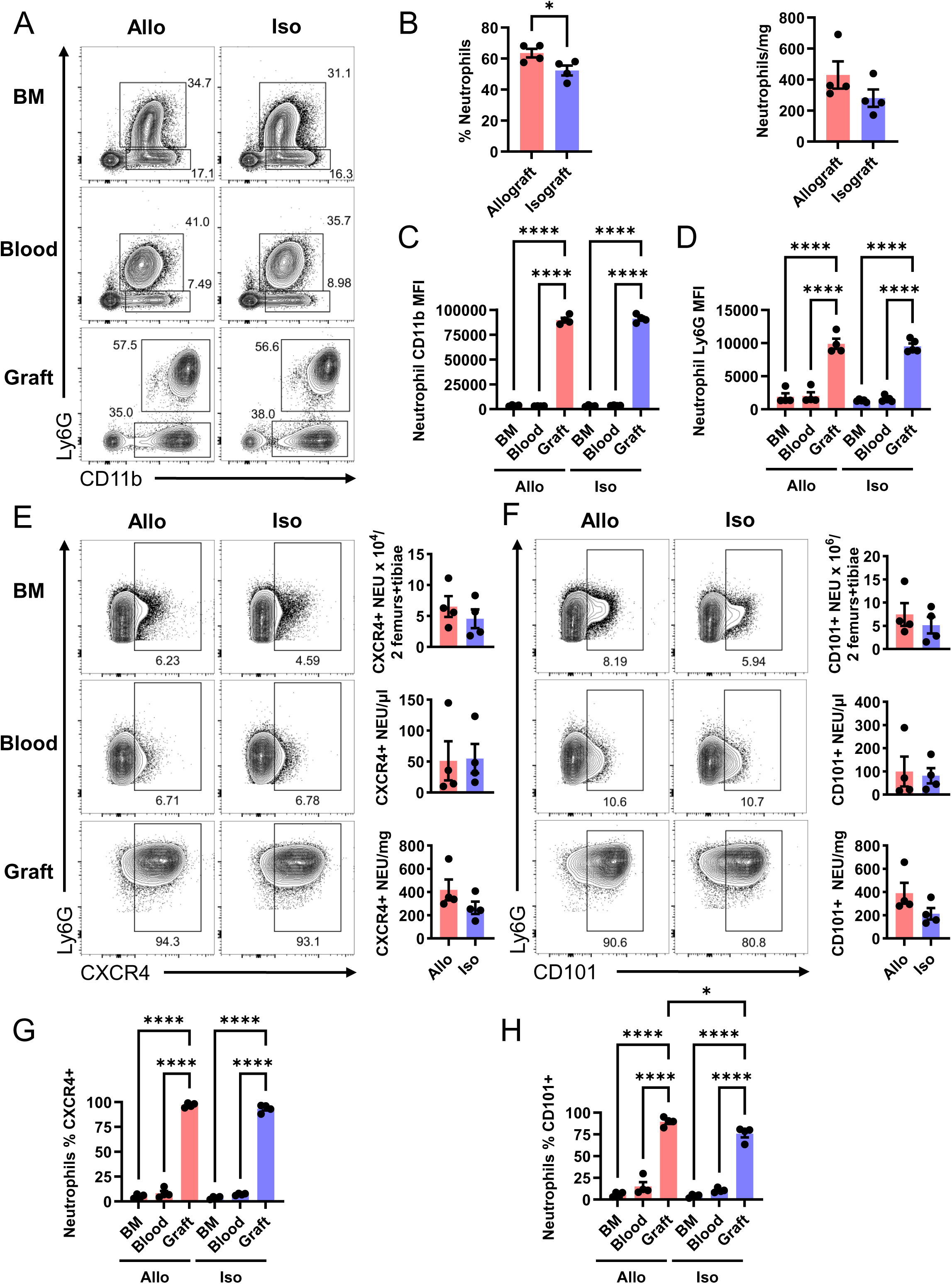
Neutrophils dominate the early graft infiltrate and intra-graft neutrophils are distinct from neutrophils elsewhere in the recipient. Groups of wild type A/J and B6 hearts were transplanted to wild type B6 recipients. Grafts, recipient blood and recipient bone marrow were harvested on day 1 post-transplant and processed for flow cytometry. **(A)** Example flow cytometry plots displaying CD11b versus Ly6G expression on live CD45+ cells from allograft and isograft digests, recipient blood and recipient bone marrow. **(B)** Frequency (of live CD45^+^ cells) and numbers (cells/mg of graft tissue) of infiltrating Ly6G^+^ neutrophils. **(C+D)** Comparison of CD11b **(C)** and Ly6G **(D)** surface expression levels on neutrophils from allograft and isograft digests, recipient blood and recipient spleen. **(E+F)** Example flow cytometry plots and summary data of cell counts from bone marrow (cells per 2 femurs + tibiae), blood (cells/μl) and grafts (cells/mg graft tissue) of CXCR4^+^ **(E)** and CD101^+^ **(F)** neutrophils. **(G+H)** Summary data comparing the percentages of neutrophils expressing CXCR4 **(G)** and CD101 **(H)** between bone marrow, blood and graft digests. Data reported as mean ± SEM, N=4 per group * = p<0.05, **** = p<0.0001

Interestingly, graft infiltrating neutrophils displayed significantly higher surface expression of CD11b and Ly6G than recipient blood and BM neutrophils (Fig 1A, C-D), indicating that graft derived neutrophils were more activated^27,28^ and mature^29^ than those in the recipient blood and BM. We confirmed this by measuring accepted indicators of neutrophil maturity (surface CD101 expression) and aging (surface CXCR4 expression). As expected, recipient blood and BM neutrophils were immature (CD101^-^) and young (CXCR4^-^), while >75% of graft infiltrating neutrophils were mature (CD101^+^) and aged (CXCR4^+^) (Fig 1E-G). This finding suggests that the IRI-induced heart graft microenvironment drives specific reprogramming of early infiltrating neutrophils.

### Neutrophils infiltrating allografts versus isografts display key phenotypic differences

As we did not find any significant differences in total numbers, maturity, or aging between neutrophils infiltrating allo- versus iso-grafts, we asked if alloimmune recognition drives differential distribution of neutrophil subsets. ISG^+^ neutrophils are rare (∼15%) in blood and BM in naïve mice.^10,30^ Flow cytometric analyses revealed a marked increase in ISG^+^ neutrophils (>70%) in the blood and BM of allo- and iso-graft recipients early after transplant (Fig 2A, C). However, we noticed a marked difference in ISG^+^ neutrophils within the heart grafts. Syngeneic isografts had five-fold lower ISG^+^ neutrophil frequency and three-fold lower absolute numbers compared to compete MHC-mismatched allografts (Fig 2A, C). In contrast, neutrophils from both allo- and iso-grafts were highly enriched for the dcTRAIL-R1^+^ protumor-angiogenic phenotype (Fig 2B, D). Unlike ISG^+^ neutrophils, dcTRAIL-R1^+^ neutrophils were minimally detectable in blood or BM from allograft and isograft recipients.

**Figure 2.**
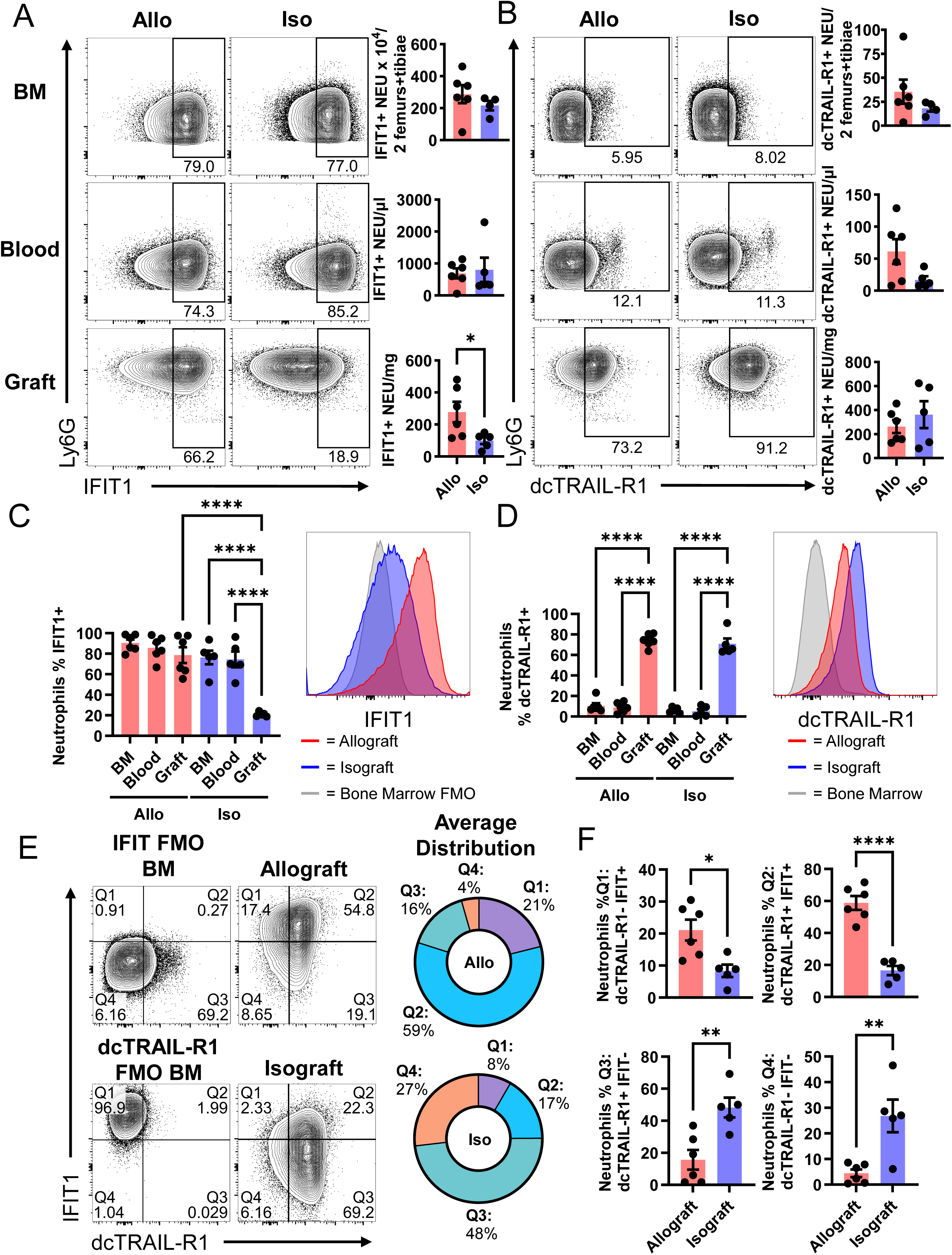
Neutrophils infiltrating allografts versus isografts display key phenotypic differences. Groups of wild type A/J and B6 hearts were transplanted to wild type B6 recipients. Grafts, recipient blood and recipient bone marrow were harvested on day 1 post-transplant. **(A-B)** Example flow cytometry plots and summary data of cell counts from bone marrow (cells per 2 femurs + tibiae), blood (cells/μl) and grafts (cells/mg graft tissue) of IFIT1^+^ **(A)** and dcTRAIL-R1^+^ **(B)** neutrophils. **(C-D)** Summary data comparing the percentages of neutrophils expressing IFIT1 **(C)** and dcTRAIL-R1 **(D)** between bone marrow, blood and graft digests. Sample histograms comparing allograft and isograft neutrophil IFIT1 **(C)** and dcTRAIL-R1 **(D)** expression. BM dcTRAIL-R1 expression and BM IFIT1 FMO are included as negative controls. **(E)** Example flow cytometry plots of IFIT1 versus dcTRAIL-R1 expression on neutrophils from allograft and isograft digests with summary average distribution of graft infiltrating neutrophil IFIT1 and dcTRAIL-R1 expression between single positive, double positive and double negative populations. FMOs were generated using bone marrow cells. **(F)** Breakdown with statistics of graft infiltrating neutrophil distribution between the four quadrants labeled in **(E)**. Data reported as mean ± SEM, N=5-6 per group * = p<0.05, ** = p<0.01, ****= p<0.0001

Although the original characterization of dcTRAIL-R1^+^ neutrophils in tumor microenvironments considered them to be transcriptomically distinct from ISG^+^ neutrophils (T3 vs T2 populations)^11^, we found a novel hybrid IFIT1^+^ dcTRAIL-R1^+^ neutrophil population representing ∼60% of neutrophils in allografts. The frequency of this hybrid neutrophil population was significantly lower in isografts (∼17%) (Fig 2E-F). While 95% of allograft infiltrating neutrophils expressed one or both of IFIT1 and dcTRAIL-R1, isografts contained a significant double negative population (∼25%) (Fig 2E-F). These findings suggest that an alloimmune response shapes phenotypic reprogramming of neutrophils following infiltration into the graft tissue.

### Recipient NK cells, but not T or B cells, drive ISG neutrophil enrichment in allografts

The clear differences in neutrophil phenotype between allo- and iso-grafts led us to ask which cellular immune component(s) mediate this differential graft specific neutrophil reprogramming. Although IFIT1 is canonically a type I interferon induced gene^31^, both type I and type II interferon can induce the ISG neutrophil phenotype.^32^ We previously showed that donor-reactive memory CD8 T cells infiltrate allografts and produce IFNγ within the first 24hr post-transplant^33^, leading us to test if a lack of early infiltrating T cell-derived IFNγ could be responsible for the lack of ISG^+^ neutrophils in isografts. To this end, we transplanted complete MHC mismatched A/J heart allografts into T and B cell deficient B6-*Rag^-/-^* recipients. Allografts transplanted to *Rag^-/-^* recipients still contained a clear ISG^+^ infiltrating neutrophil population (∼35%) (Fig 3A-C).

**Figure 3.**
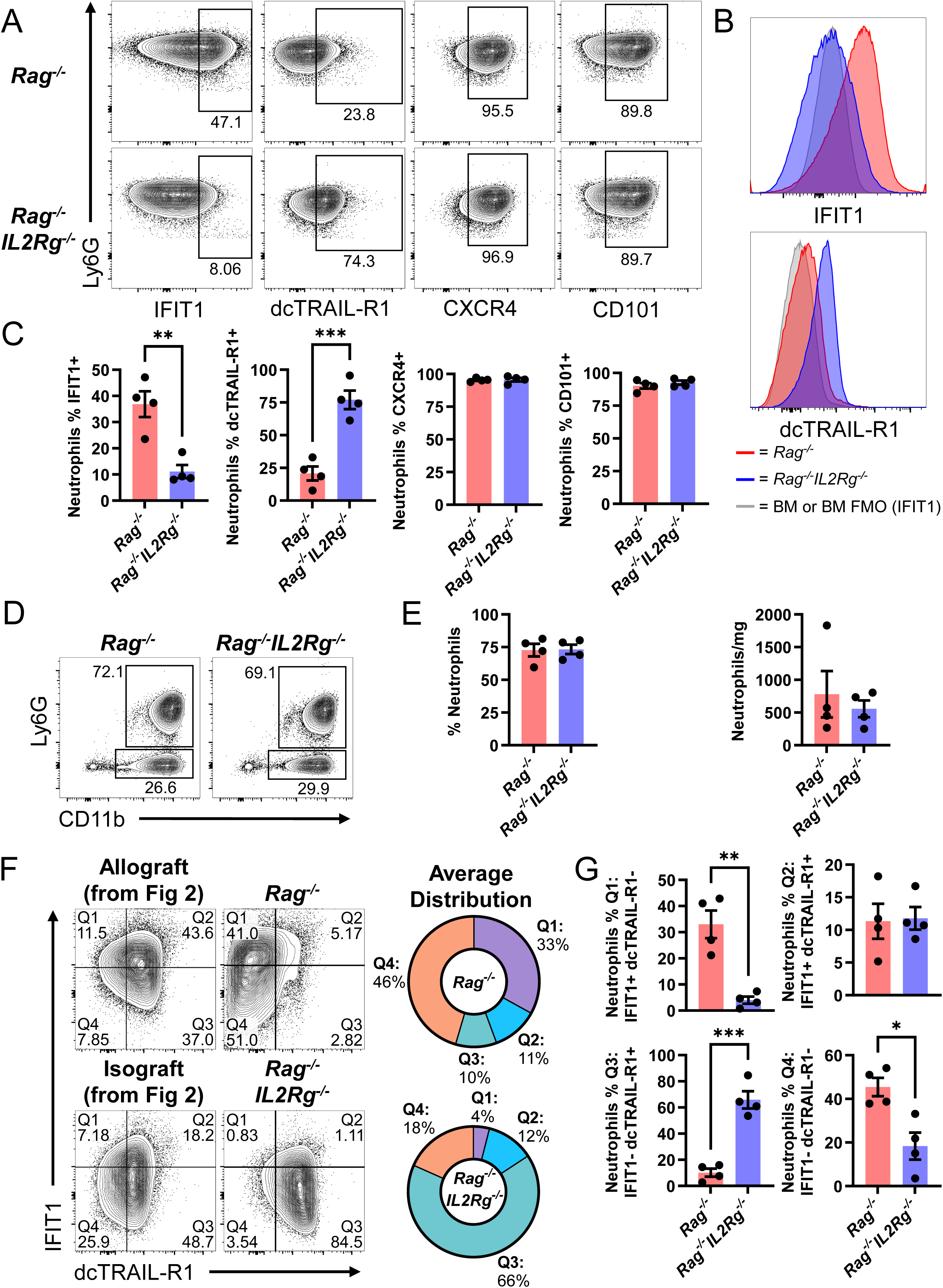
Recipient NK cells, but not T or B cells, mediate ISG neutrophil enrichment in allografts. Groups of wild type A/J hearts were transplanted to *Rag^-/-^* and *Rag^-/-^IL2Rg^-/-^* B6 recipients. Grafts, recipient blood and recipient bone marrow were harvested on day 1 post-transplant and processed for flow cytometry. **(A)** Example flow cytometry plots of neutrophils expressing IFIT1, dcTRAIL-R1, CXCR4 and CD101 in allografts transplanted to *Rag^-/-^* and *Rag^-/-^IL2Rg^-/-^* recipients. **(B)** Sample histograms comparing allograft neutrophil IFIT1 and dcTRAIL-R1 expression between *Rag^-/-^* and *Rag^-/-^IL2Rg^-/-^* recipients. BM dcTRAIL-R1 expression and BM IFIT1 FMO are included as negative controls. **(C)** Summary expression data comparing the frequencies of IFIT1, dcTRAIL-R1, CXCR4 and CD101 expressing neutrophils in allografts transplanted to *Rag^-/-^* and *Rag^-/-^IL2Rg^-/-^* recipients. (**D)** Example flow cytometry plots of CD11b versus Ly6G expression on live CD45+ cells from allografts transplanted to *Rag^-/-^* and *Rag^-/-^IL2Rg^-/-^* recipients. **(E)** Summary data comparing the percent Ly6G^+^ neutrophils (of live CD45+ cells) and numbers of neutrophils per mg of graft tissue between allografts transplanted to *Rag^-/-^* and *Rag^-/-^IL2Rg^-/-^* recipients. **(F)** Example flow cytometry plots of IFIT1 versus dcTRAIL-R1 expression on neutrophils from allografts transplanted to *Rag^-/-^* and *Rag^-/-^IL2Rg^-/-^* recipients with summary average distribution of graft infiltrating neutrophil IFIT1 and dcTRAIL-R1 expression between single positive, double positive and double negative populations. Additional plots of allograft and isograft samples are included as examples for comparison. **(G)** Breakdown with statistics of graft infiltrating neutrophil distribution between the four quadrants labeled in **(F)**. Data reported as mean ± SEM, N=4 per group * = p<0.05, ** = p<0.01, *** = p<0.001, ****=p<0.0001

The limited impact of recipient T or B cells on maintenance of the ISG^+^ neutrophil program after infiltrating an allograft led us to consider the possibility of innate allorecognition by allograft infiltrating NK cells as an alternative explanation for the difference in ISG^+^ neutrophil frequency between allo and isografts. To test the impact of NK cells on the maintenance of the ISG^+^ neutrophil phenotype after infiltrating an allograft, we compared allografts transplanted to *Rag^-/-^* versus *Rag^-/-^IL2Rg^-/-^* (lacking T, B and NK cells) recipients. There was no difference in the frequency (of live CD45^+^ cells) or number of neutrophils (per mg of graft tissue) between allografts harvested from *Rag^-/-^* versus *Rag^-/-^IL2Rg^-/-^* recipients (Fig 3D-E).

Interestingly, the frequency of ISG^+^ neutrophils was dramatically reduced (∼3-fold) in allografts from *Rag^-/-^IL2Rg^-/-^* compared to *Rag^-/-^* recipients (Fig 3A-C), to approximately the level found in isografts from WT recipients (Fig 2A). However, the frequency of ISG^+^ neutrophils in the BM of *Rag^-/-^IL2Rg^-/-^* recipients was significantly reduced compared to the BM of *Rag^-/-^* recipients, indicating that NK cells may also influence initial ISG neutrophil programming in the BM (Fig S3). Strikingly, loss of NK cells (*Rag^-/-^IL2Rg^-/-^* recipients) reversed the lower proportion of dcTRAIL-R1^+^ neutrophils seen in *Rag^-/-^* allografts to a level similar to that seen in allo- and iso-grafts from WT recipients (Fig, 2E, Fig 3A-C). The concurrent reduction in ISG^+^ and increase in dcTRAIL-R1^+^ neutrophils in allografts from *Rag^-/-^IL2Rg^-/-^* recipients may indicate shared but opposing regulatory mechanisms of inflammatory ISG (marked by IFIT1 expression) versus proangiogenic (marked by dcTRAIL-R1 expression) phenotypic programming. The pattern of IFIT1 versus dcTRAIL-R1 expression on neutrophils from *Rag^-/-^* versus *Rag^-/-^IL2Rg^-/-^* recipient allografts was characterized by a shift between the two single positive populations (decreased IFIT1^+^dcTRAIL-R1^-^ and increased IFIT1-dcTRAIL-R1^+^ cells) with no change in the proportion of double positive cells (Fig 3E-F). This contrasts with allograft infiltrating neutrophils from WT recipients, where 60% of neutrophils co-expressed IFIT1 and dcTRAIL-R1 (Fig 2E-F, 3E).

Like those in allo- and iso-grafts from WT recipients, neutrophils in allografts from *Rag^-/-^* and *Rag^-/-^IL2Rg^-/-^* recipients were predominantly aged/CXCR4^+^ and mature/CD101^+^ cells (Fig 3A-C). *Rag^-/-^* and *Rag^-/-^IL2Rg^-/-^* recipient BM and blood neutrophils matched the mostly CXCR4^-^ CD101^-^ profile observed in WT recipients (Fig S3). The pattern of CD11b and Ly6G expression on neutrophils in *Rag^-/-^* and *Rag^-/-^IL2Rg^-/-^* recipients (high on graft neutrophils and low on BM and blood neutrophils) also matched that seen in WT allo- and iso-graft recipients (Fig S3).

### Genetic deficiency of lymphocyte populations in allograft recipients impacts graft ISG expression

To better understand the allograft environment in these conditions with differing levels of ISG neutrophils, we performed a transcriptomic analysis of bulk RNA from allografts that were transplanted to wild type, *Rag^-/-^* and *Rag^-/-^IL2Rg^-/-^* recipients using NanoString nCounter analysis (770-gene PanCancer Immune Profiling codeset) (Fig 4 A-F).

**Figure 4.**
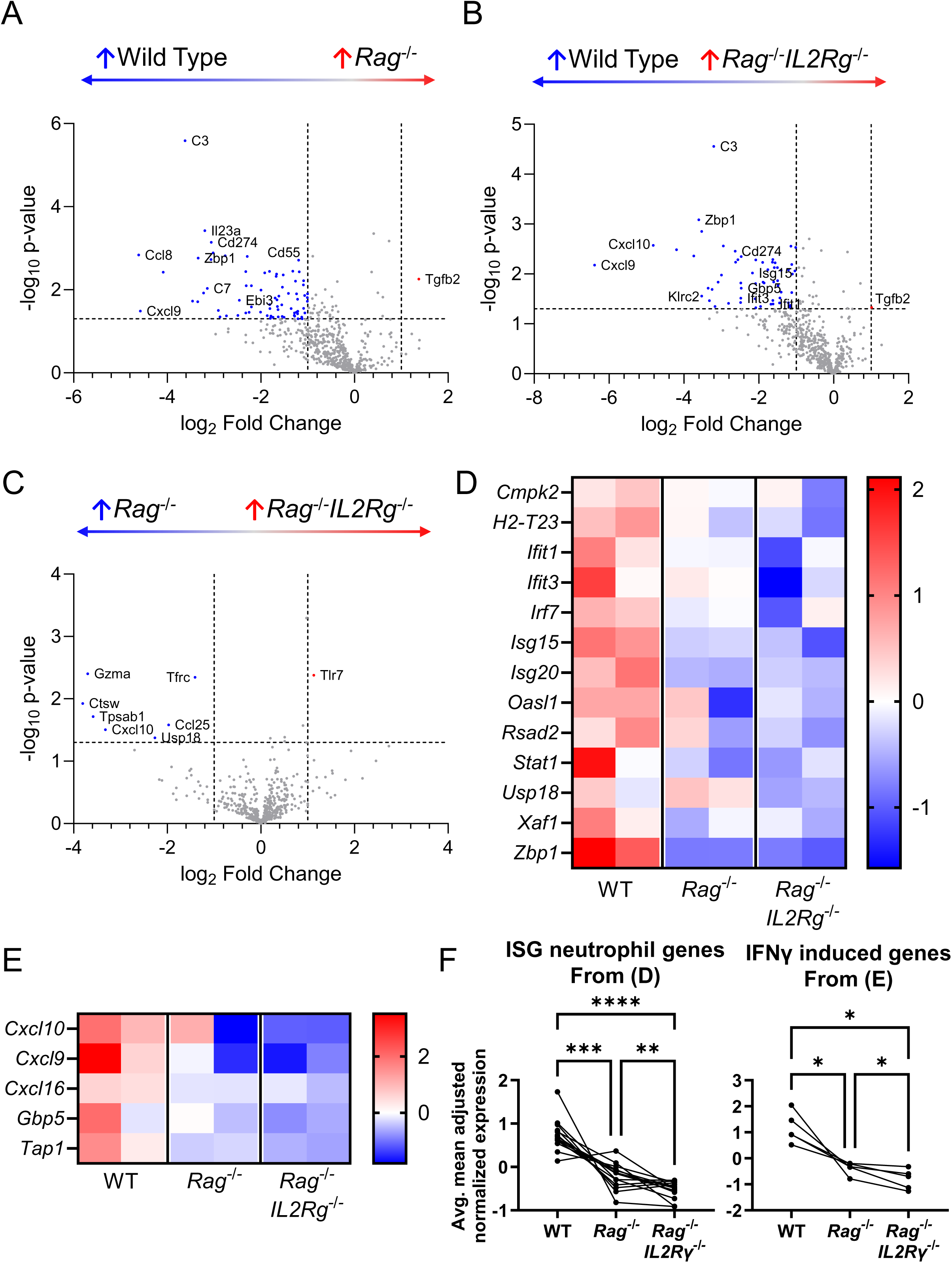
Differential IFN response expression signature between allografts transplanted to WT, *Rag^-/-^* and *Rag^-/-^IL2Rg^-/-^* recipients. Groups of wild type A/J hearts were transplanted to wild type, *Rag^-/-^* and *Rag^-/-^IL2Rg^-/-^* B6 recipients and harvested on day 1 post-transplant. NanoString nCounter RNA expression analysis was performed using the mouse PanCancer Immune Profiling panel on RNA isolated from bulk graft tissue homogenates (n=2 per group). **(A-C)** Volcano plots showing the distribution of DEGs between allografts from WT versus *Rag^-/-^IL2Rg^-/-^* recipients **(A)**, WT versus *Rag^-/-^* recipients **(B)**, and *Rag^-/-^* and *Rag^-/-^IL2Rg^-/-^* (C) recipients. Dashed lines indicate cutoffs of ± 2-fold change and p<0.05. Heatmaps listing all DEGs for each comparison can be found in **Supplemental Figure 3**. **(D)** Heatmap showing the relative expression in each sample of 13 G5b/ISG neutrophil signature genes (from Xie et al., 2020)^10^ in the PanCancer Immune Profiling Panel. **(E)** Heatmap showing the relative expression in each sample of selected IFNγ induced genes. **(F)** Averaged mean adjusted expression levels in each group for each of the 13 G5b/ISG signature genes (left) and each of the selected IFNγ induced genes (right). Statistical comparisons in **(F)** were calculated using repeated measures one-way ANOVAs with Geisser-Greenhouse correction and Šídák’s multiple comparisons test. N=2 transplants per group. * = p<0.05, ** = p<0.01, *** = p<0.001, ****= p<0.0001

Over 50% of differentially expressed genes (DEGs) between allografts from WT versus *Rag^-/-^* and WT versus *Rag^-/-^IL2Rg^-/-^* recipients were shared (Fig 4A-B, Fig S4-S5). Enrichr pathway analysis^23–25^ of the set of shared DEGs using the Reactome pathway database^26^ notably included IFNα/β (8 genes) and IFNγ (6 genes) signaling among the top ten enriched pathways. Also prominent among the shared DEGs were several complement components. Interestingly, pathway analysis on the unique DEGs from the two comparisons found multiple pathways related to interferon signaling present only in the WT versus *Rag^-/-^IL2Rg^-/-^* specific genes. A broader decrease in interferon signaling in the allografts from *Rag^-/-^IL2Rg^-/-^* versus *Rag^-/-^* recipients supports our hypothesis that reduced interferon levels in these grafts leads to the loss of ISG expression in infiltrating neutrophils.

Direct comparison of allografts from *Rag^-/-^* and *Rag^-/-^IL2Rg^-/-^* recipients yielded a limited number of differentially expressed genes (DEGs), the most notable of which was CXCL10, a chemokine that is a highly sensitive marker of IFNγ response (Fig 4C). Lower CXCL10 expression in allografts from *Rag^-/-^IL2Rg^-/-^* recipients further supports the hypothesis that the reduced IFNγ signaling in *Rag^-/-^IL2Rg^-/-^* recipients, contributes to the lower frequency of ISG^+^ neutrophils in these grafts. There was a smaller but also significant decrease in expression of *Usp18*, which can be induced by both type I and type III IFNs, in allografts from *Rag^-/-^IL2Rg^-/-^* recipients. Allografts from *Rag^-/-^IL2Rg^-/-^* recipients also had lower expression of the cytotoxic granule components cathepsin W and granzyme A, in line with the loss of recipient NK cells.

The NanoString panel contained 13 out of 30 ISG^+^ neutrophil signature genes.^10^ Although interpretation is complicated by the use of RNA from bulk graft tissue homogenate, when compared to allografts from WT recipients, 4/13 ISG neutrophil signature genes had significantly lower expression in allografts from *Rag^-/-^* recipients (Fig 4A, S4A) versus 8/13 that were significantly lower in allografts from *Rag^-/-^IL2Rg^-/-^* recipients (Fig 4B, S4B). Mean adjusted average expression levels of the ISG^+^ neutrophil signature genes showed a clear stepwise downward trend in allografts from WT to *Rag^-/-^* to *Rag^-/-^IL2Rg^-/-^* recipients, matching the stepwise decrease in ISG^+^ neutrophils in allografts from WT to *Rag^-/-^* to *Rag^-/-^IL2Rg^-/-^* recipients (Fig 4D, F). Similar analysis of the expression levels of a set of IFNγ induced genes in the NanoString panel showed the same stepwise downward trend in allografts from WT to *Rag^-/-^* to *Rag^-/-^IL2Rg^-/-^* recipients, supporting our hypothesis that diminished intra-graft IFNγ signaling may be responsible for down modulating the ISG program in neutrophils after entering the graft (Fig 4E, F).

### NK cells secrete IFNγ when infiltrating allo-but not iso-grafts

We then compared the NK cell phenotypes in WT allograft and isograft recipients to explain the NK cell-dependent allograft specific neutrophil phenotypic programming. There was no significant difference in the frequency or numbers of total NK cells in allografts versus isografts harvested from WT recipients (Fig S1). Both allograft and isograft infiltrating NK cells were predominantly immature (NK1.1^+^ DX5^-^ NKp46^-^)^34,35^, which was also the dominant phenotype of NK cells in the recipient BM (Fig 5A, B). This contrasted with recipient spleen derived NK cells which were approximately 50% mature (NK1.1^+^ DX5^+^ NKp46^+^). There was also no significant difference in the absolute numbers of mature versus immature graft infiltrating NK cells between allo- and iso-grafts (Fig 5C). Graft infiltrating NK cells had significantly higher cell surface expression of the degranulation marker LAMP-1/CD107a than infiltrating T cells, although there was no difference between allograft and isograft infiltrating NK cells (Fig 5D-E). Recipient spleen and bone marrow NK cells had much lower cell surface CD107a levels, approximating those of the graft infiltrating T cells (Fig S6).

**Figure 5.**
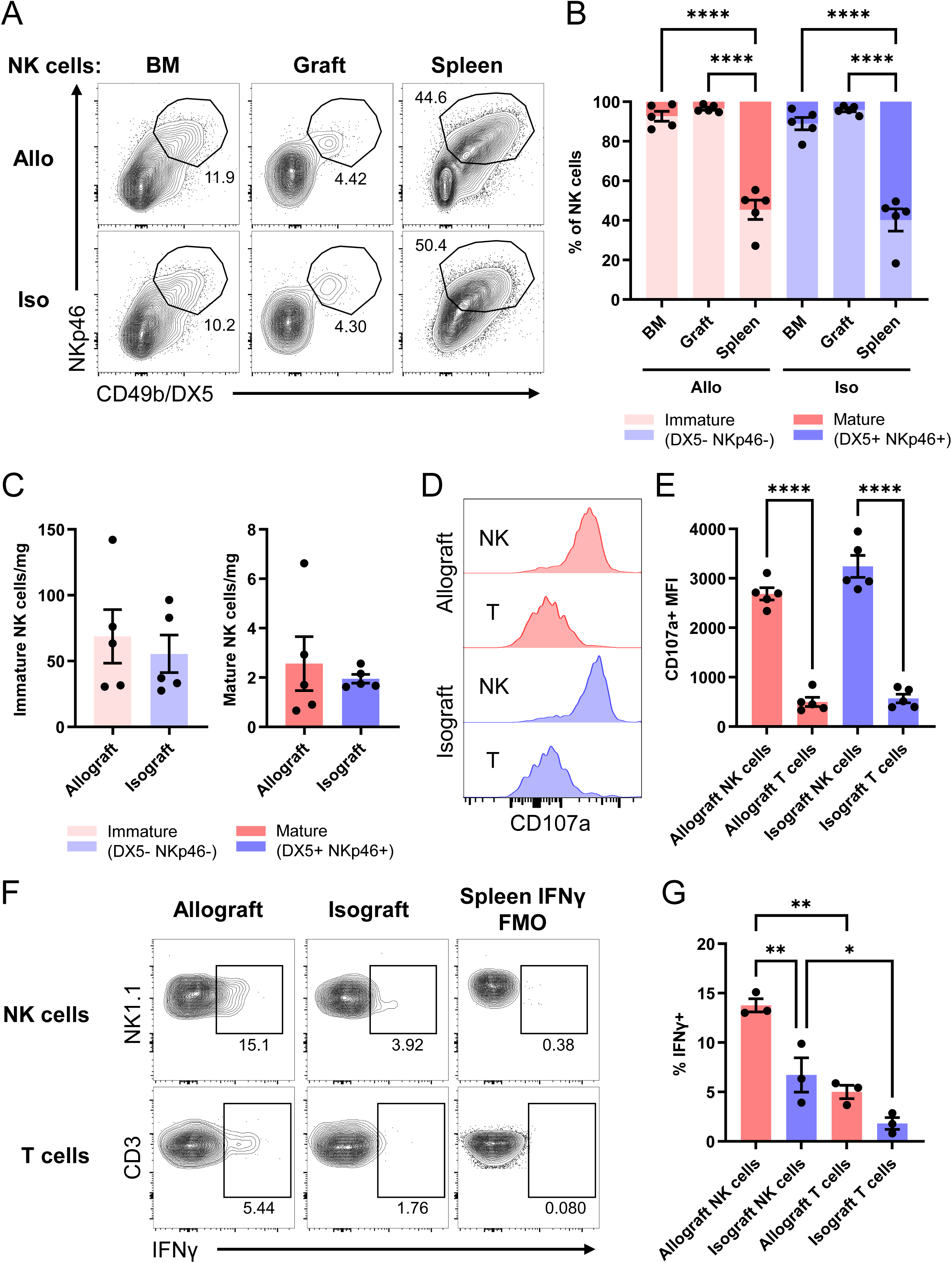
NK cells secrete IFNγ when infiltrating allo-but not iso-grafts. Groups of wild type A/J and B6 hearts were transplanted to wild type B6 recipients. Grafts, recipient bone marrow and recipient spleens were harvested on day 1 post-transplant. **(A)** Example flow cytometry plots showing CD49b/DX5 versus NKp46 expression in allograft and isograft infiltrating, recipient bone marrow and recipient spleen CD3^-^ NK1.1^+^ NK cells. **(B)** Breakdown of mature (CD49b/DX5^+^ NKp46^+^) versus immature (CD49b/DX5^-^NKp46^-^) NK cells in each site from **(A)** in allograft and isograft recipients. **(C)** Comparison of numbers per mg graft tissue of mature and immature NK cells between allografts and isografts. **(D-E)** Example histograms **(D)** and summary data **(E)** showing the distribution of CD107a/LAMP1 expression on allograft and isograft infiltrating CD3^+^ T cells and CD3^-^ NK1.1^+^ NK cells. N=5 per group. **(F-G)** Bulk cells from digested allo- and iso-grafts (N=3 per group) were cultured ex vivo with PMA, ionomycin, monensin and brefeldin A for 4hr before fixation and permeabilization for intracellular flow cytometry staining. **(F)** Sample flow cytometry plots and **(G)** summary data with statistics showing the proportion of IFNγ^+^ T cells and NK cells in allografts and isografts after culture. FMOs were generated using cultured bulk spleen cells. Data reported as mean ± SEM, * = p<0.05, ** = p<0.01, *** = p<0.001, ****= p<0.0001

Because NK cell maturation is associated with shifts in the expression of cytotoxic and cytokine producing functions^36^, we tested graft infiltrating NK and T cell production of cytokines and cytotoxic mediators following stimulation with PMA and ionomycin. In line with our hypothesis, we found a significantly higher frequency of IFNγ producing graft infiltrating NK cells in allografts compared to isografts (Fig 5E-F). In contrast, the frequency of IFNγ producing graft infiltrating T cells after stimulation was significantly lower than the frequency of IFNγ producing NK cells in both allo- and iso-grafts (Fig 5E-F). Graft infiltrating NK cells also exhibited higher levels of intracellular perforin and granzyme B expression than graft infiltrating T cells, but, in contrast to IFNγ, there were no differences between allo- and iso-grafts (Fig S6).

These findings suggest that graft infiltrating NK cells have both greater numbers and higher expression of effector functions than infiltrating T cells at this timepoint, reinforcing the findings from *Rag^-/-^* and *Rag^-/-^IL2Rg^-/-^* mice showing a pivotal role for IFNγ-producing NK cells in shaping the phenotypic programing of early graft infiltrating neutrophils.

## Discussion

Using the mouse heterotopic heart transplant model, we show for the first time that the graft environment and alloimmune response combine to reprogram infiltrating neutrophils to exhibit a distinct phenotype. Upon entering an allogeneic or syngeneic heart graft, neutrophils rapidly mature, age and acquire a proangiogenic phenotype marked by dcTRAIL-R1 expression. In contrast, we found a differential impact of donor-recipient MHC mismatch on the maintenance of ISG expression by infiltrating neutrophils. While the ISG signature was sustained in allograft infiltrating neutrophils, it was mostly turned off in isograft infiltrating neutrophils. This difference is notable given our finding that post-surgical inflammation induced rapid expansion of immature ISG^+^ neutrophils in the recipient BM and peripheral blood of both allograft and isograft recipients. Mechanistically, studies using *Rag^-/-^* and *Rag^-/-^IL2Rg^-/-^* allograft recipients indicated that NK cell allorecognition sustains ISG expression by early allograft infiltrating neutrophils. Further supporting the role of NK cells, graft infiltrating NK cells are more abundant than T or B cells at this timepoint and allograft infiltrating NK cells show enhanced IFNγ production compared to infiltrating T cells or isograft infiltrating NK cells.

The lack of dcTRAIL-R1 on blood and BM neutrophils and acquisition of dcTRAIL-R1 expression upon entering the heart graft matches the initial report describing dcTRAIL-R1 expressing neutrophils, which only identified these cells in tumor tissue where they were enriched in more hypoxic areas of tumors.^11^ Hypoxia is also prominent in injured tissues early after an ischemic insult, which supports our observation that surface dcTRAIL-R1 expression was increased on neutrophils upon entering the hypoxic heart graft microenvironment. Whether this hypoxic environment also causes the neutrophil maturation and aging seen upon entering a heart graft is unclear. Neutrophil maturation and aging are associated with greater expression of effector functions associated with inflammation and tissue injury^7,8^, such as occurs during IRI.

Inflammation induces expansion of ISG^+^ neutrophils in the bone marrow.^10^ Herein, we observed similar expansion of ISG^+^ neutrophils in the bone marrow of both allo- and iso-graft recipients which was likely induced by systemic inflammation from the surgery. However, upon entering the heart graft, maintenance of the ISG^+^ program was dependent on donor-recipient MHC mismatch. In an allograft the BM encoded inflammatory ISG program was maintained alongside the microenvironment imposed dcTRAIL-R1^+^ proangiogenic program, generating a dominant hybrid IFIT1^+^ dcTRAIL-R1^+^ neutrophil population. In the absence of allorecognition, the heart graft microenvironment turned off the inflammatory ISG program in infiltrating neutrophils. Interestingly, isografts and allografts transplanted to *Rag^-/-^* recipients contained a significant proportion of IFIT^-^ dcTRAIL-R1^-^ neutrophils, indicating a need for future studies that more deeply characterize graft infiltrating neutrophils in these and other transplant conditions.

Despite our clear findings, there are limitations to studies using the murine heterotopic heart transplant model. Heart grafts in this model are non-life sustaining and mechanically unloaded, which impacts heart tissue remodeling after reperfusion and does not match the clinical use of heart transplantation. Importantly, our study uses healthy murine heart transplant recipients, whereas clinical heart transplant recipients are patients with end-stage heart disease having systemic inflammation and other related comorbidities at the time of transplant. The pre-transplant presence of underlying inflammation in human heart transplant patients may mean that they already have augmented generation of ISG^+^ neutrophils in the bone marrow compared to healthy control patients and our mouse heart transplant recipients, but this remains to be determined. Whether preexisting inflammation impacts neutrophil reprogramming after infiltrating a heart graft is unknown at this time. Additionally, we do not assess the kinetics of these differences in neutrophil reprogramming, instead choosing to focus on a single 24 hr post-reperfusion timepoint based on work from our group^16^ and others^17^ indicating that this time is the peak of early neutrophil infiltration and activation during heart transplant IRI.

In summary, our findings provide novel insights into the regulation of key neutrophil subsets previously described in infection and cancer. This work also represents the first attempt to understand how alloimmune responses and the unique microenvironment of a solid organ transplant contribute to neutrophil heterogeneity and plasticity. Given the previously established important role of neutrophils in the early immune response to a solid organ transplant, these findings may enable more selective targeting of detrimental neutrophil subsets to enhance solid organ transplant outcomes.

Although, our results indicate that NK cell derived IFNγ helps maintain the ISG program in allograft infiltrating neutrophils, simply blocking IFNγ is not a viable therapeutic strategy in transplantation. Studies in animal models raise clear concerns about peri-transplant anti-IFNγ treatment due to its pleiotropic effects in the early post-reperfusion environment.^37–39^ Thus, deeper profiling is needed to identify alternative targets that can either eliminate specific subpopulations of graft infiltrating neutrophils or shift their phenotype away from the ISG-signature, optimally towards more beneficial functions that promote tissue repair. Expansion of the ISG neutrophil population has been associated with disease severity in infections (malaria^40^ and tuberculosis^41^) and autoimmune diseases (lupus^42^), and with cancer responsiveness to immune checkpoint inhibitor therapy.^32^ Novel insights into regulation of the ISG neutrophil phenotype gleaned from studying these cells in transplantation could be widely beneficial across a variety of inflammatory and disease states.

## Novelty and Significance

### What is known?

- Recent developments in neutrophil biology have revealed substantial phenotypic and functional neutrophil heterogeneity in homeostasis, malignancy and infection.
- Neutrophils are highly plastic, with their cell states differentially modulated upon entering the microenvironments of different organs including the heart.
- Little is known about neutrophil heterogeneity in solid organ transplants, including whether and how ischemia-reperfusion injury couples with donor-reactive immune responses to influence neutrophil reprograming after infiltrating a heart graft.

### What new information does this article contribute?

- We reveal for the first time that alloimmune responses shape neutrophil reprogramming during infiltration into a transplanted heart, with neutrophils in a heart allograft maintaining an inflammatory interferon stimulated gene (ISG) expression program that is dependent on NK cell activation, while neutrophils in an isograft turn off this expression.
- The heart graft microenvironment drives additional specific reprogramming of early infiltrating neutrophils towards a mature, aged and pro-angiogenic dcTRAIL-R1^+^ phenotype regardless of the degree of donor-recipient mismatch.
- In an allograft, neutrophils maintain the bone marrow encoded inflammatory ISG program alongside the heart microenvironment imposed dcTRAIL-R1^+^ proangiogenic program, generating a dominant, previously undescribed, hybrid IFIT1^+^ dcTRAIL-R1^+^ population.

Our findings provide novel insights into the regulation and plasticity of key neutrophil subsets previously described in infection and cancer. This work also indicates how alloimmune responses and the unique microenvironment of a heart transplant contribute to neutrophil heterogeneity and plasticity. Given the previously established important role of neutrophils in the early immune response to a solid organ transplant, these findings should facilitate more selective targeting of detrimental neutrophil subsets, such as ISG^+^ neutrophils, to improve heart transplant outcomes. Understanding how neutrophil ISG programming is silenced after infiltrating a syngeneic heart transplant could also provide insights into other states of neutrophilic acute heart inflammation, including myocardial infarction and myocarditis.

## Acknowledgements

The authors thank Drs. William Baldwin (Cleveland Clinic), Neil Greenspan (University Hospitals and Case Western Reserve University School of Medicine), Alan Levine (Case Western Reserve University School of Medicine) and Anna Valujskikh (Cleveland Clinic) for their valuable advice throughout the development and undertaking of this project.

