## Supplementary Materials for "NK cell allorecognition shapes reprogramming of neutrophils infiltrating heart allografts"

**Supplemental Materials**

Figures S1-6

Major Resources Table

### SUPPLEMENTAL FIGURE 1

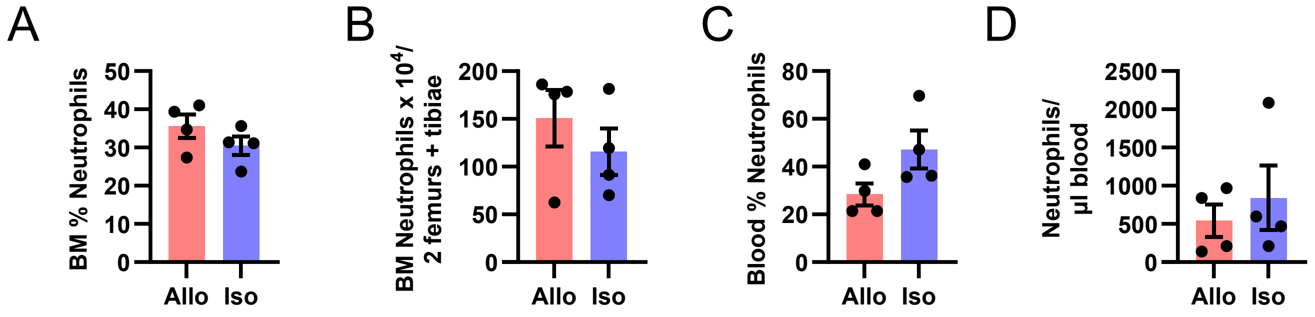

**Supplemental Figure 1. Bone marrow and blood neutrophil frequencies and numbers from allo- and iso-graft recipients.** Additional data related to Fig 1. Groups of wild type A/J and B6 hearts were transplanted to wild type B6 recipients. Recipient BM and blood were harvested on day 1 post-transplant and processed for flow cytometry to enumerate $CD11b^+$ $Ly6G^+$ neutrophils. Summary data comparing neutrophil frequency (of total live $CD45^+$ cells) **(A, C)** numbers per 2 femurs + tibiae (BM) **(B)** or µl blood **(D)** between allograft and isograft recipients. Data reported as mean ± SEM, N=4 per group.

### SUPPLEMENTAL FIGURE 2

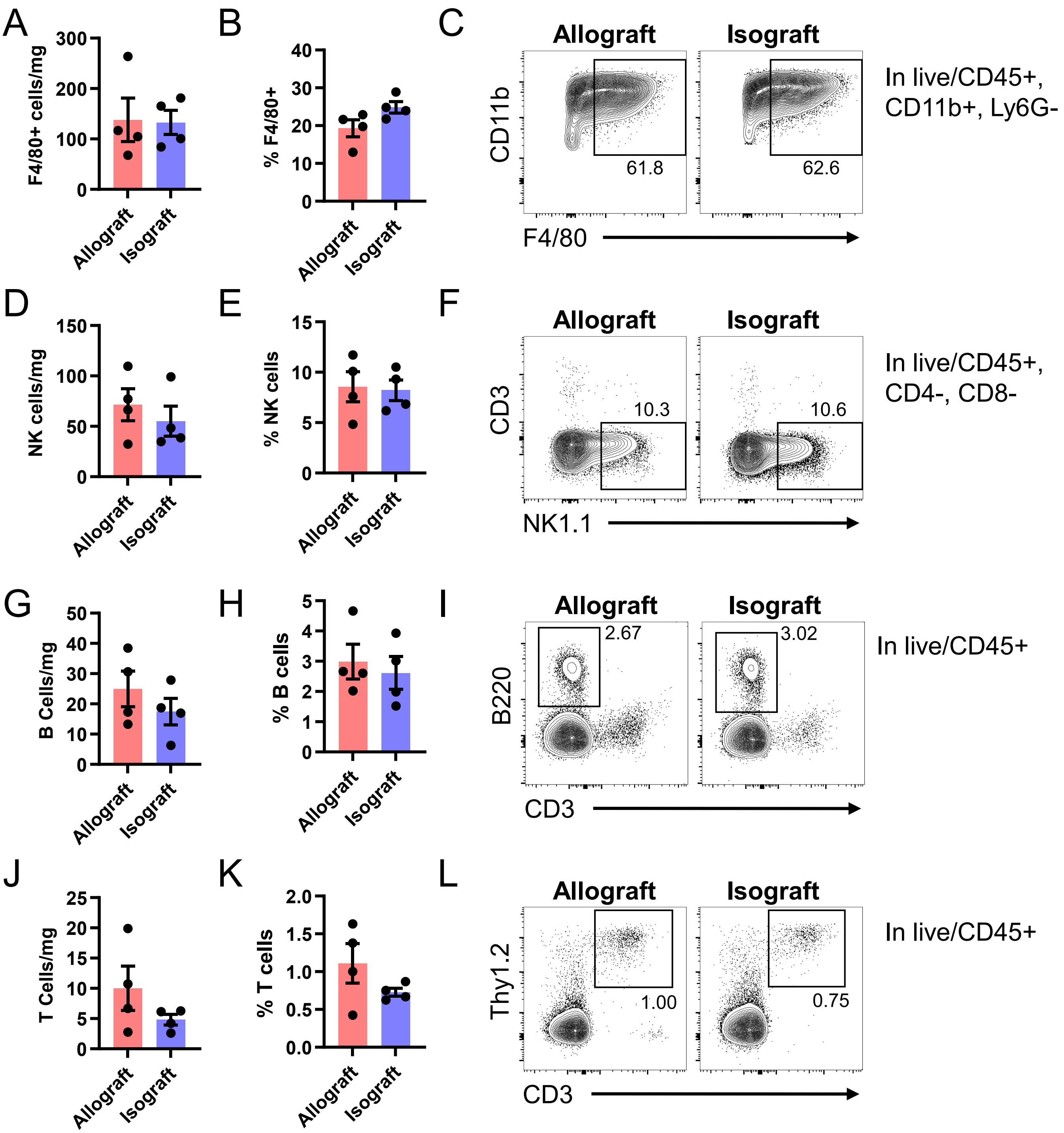

**Supplemental Figure 2. Numbers and frequencies of non-neutrophil immune cell populations in allo- and iso-grafts.** Additional data related to **Fig 1A-B**. Groups of wild type A/J and B6 hearts were transplanted to wild type B6 recipients. Grafts were harvested on day 1 post-transplant and processed for flow cytometry to enumerate F4/80+ macrophages **(A-C)**, $CD3^-$ $NK1.1^+$ NK cells **(D-F)**, $B220^+$ B cells **(G-I)** and $Thy1^+$ $CD3^+$ T cells **(J-L)**. Example flow cytometry plots identifying different graft immune cell populations. Summary data comparing numbers of cells per mg of graft tissue and the proportion of live $CD45^+$ cells for each population between allografts and isografts. Data reported as mean ± SEM, N=4 per group

### SUPPLEMENTAL FIGURE 3

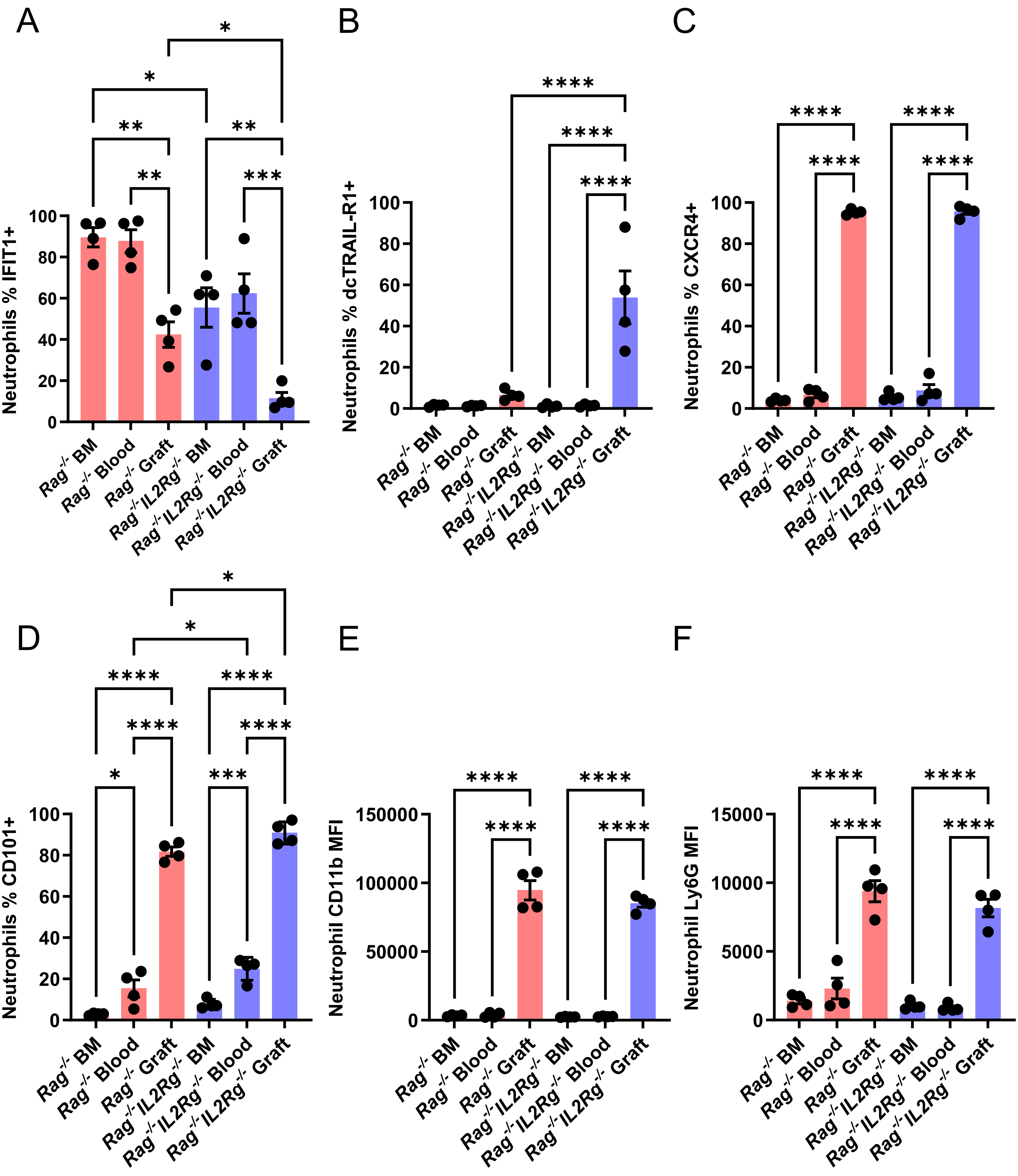

**Supplemental Figure 3. Comparison of surface expression of various surface markers on neutrophils from allografts, recipient blood, and recipient bone marrow from *Rag$^{-/-}$* and *Rag$^{-/-}$IL2Rg$^{-/-}$* recipients.** Additional data related to **Fig 3**. Groups of wild type A/J hearts were transplanted to *Rag$^{-/-}$* and *Rag$^{-/-}$IL2Rg$^{-/-}$* B6 recipients. Grafts were harvested on day 1 post-transplant and processed for flow cytometry. Summary data comparing **(A-D)** the percentages of neutrophils expressing IFIT1 **(A)**, dcTRAIL-R1 **(B)**, CXCR4 **(C)**, and CD101 **(D),** and **(E-F)** the neutrophil surface expression of CD11b **(E)** and Ly6G **(F)** between bone marrow, blood and graft digests. Data reported as mean ± SEM, N=4 per group. ✱ = $p<0.05$, ✱✱ = $p<0.01$, ✱✱✱ = $p<0.001$, ✱✱✱✱= $p<0.0001$

### SUPPLEMENTAL FIGURE 4

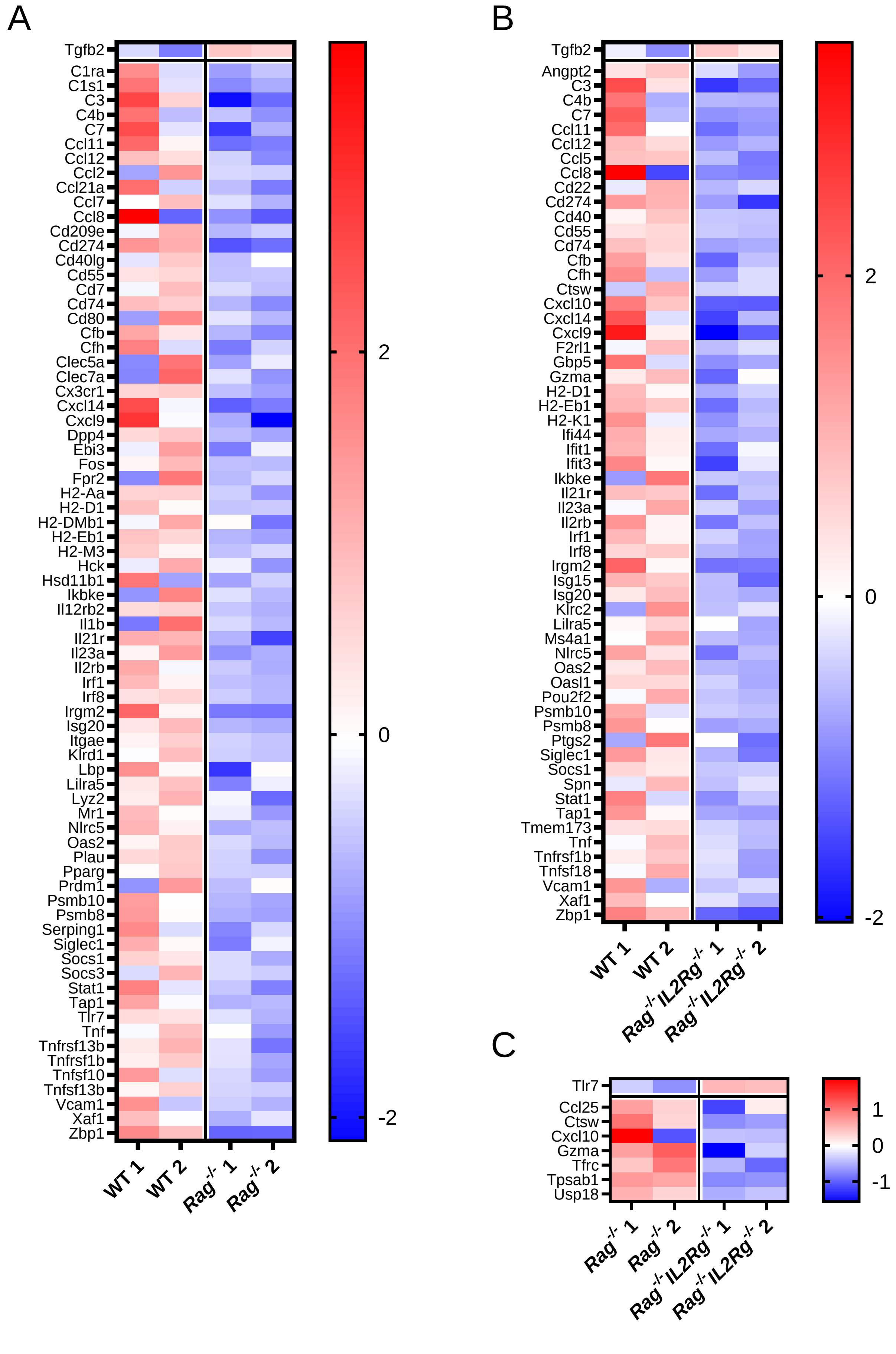

**Supplemental Figure 4. Heatmaps of full DEG lists from NanoString comparison of allografts from WT vs *Rag*$^{-/-}$ (A), WT vs *Rag*$^{-/-}$*IL2Rg*$^{-/-}$ (B), and *Rag*$^{-/-}$ vs *Rag*$^{-/-}$*IL2Rg*$^{-/-}$ (C) recipients.** Additional data corresponding to Fig 4A-C.

### SUPPLEMENTAL FIGURE 5

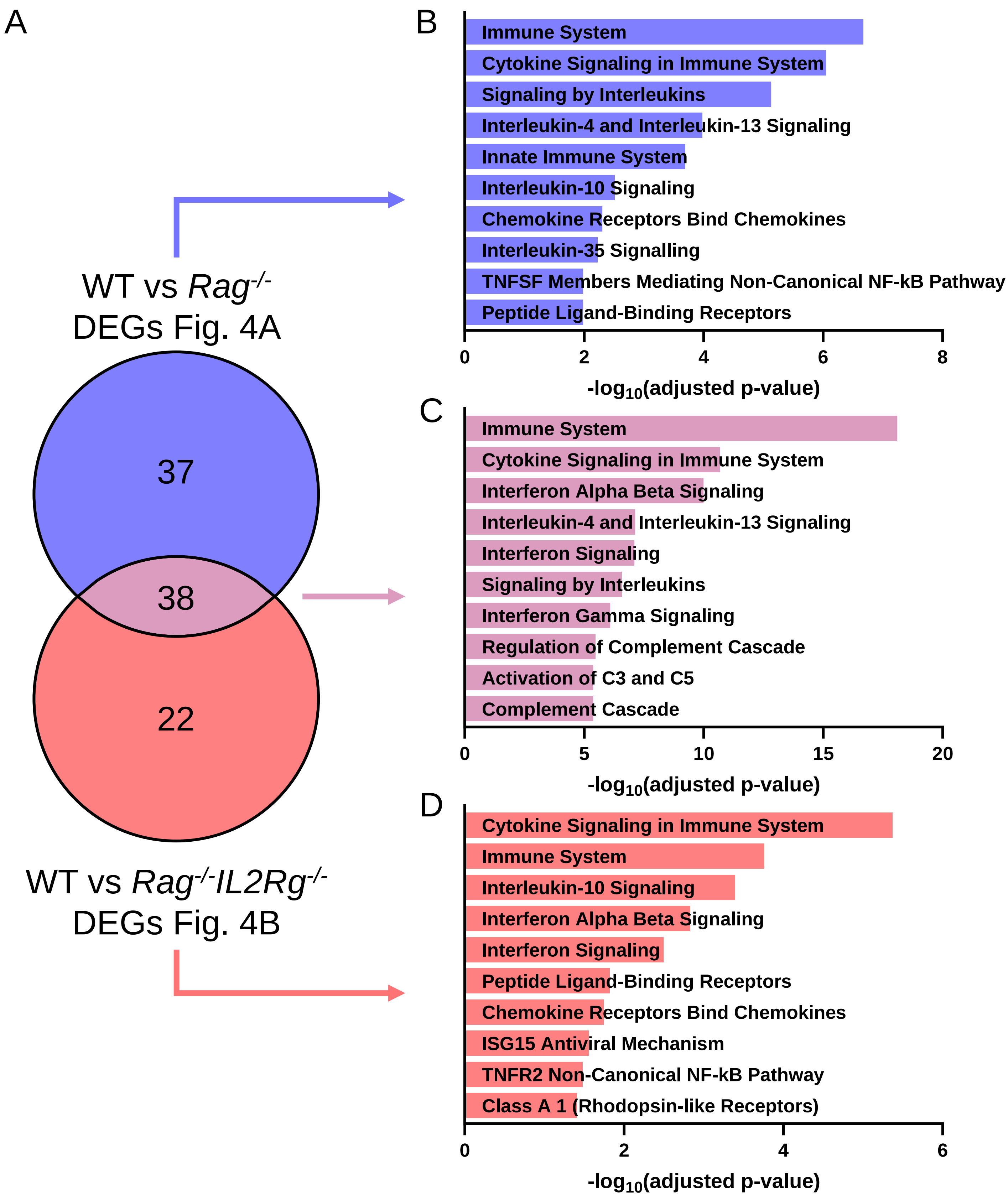

**Supplemental Figure 5. Enrichr pathway analysis of unique and shared DEGs from NanoString analysis of allografts from WT vs *Rag$^{-/-}$* and WT vs *Rag$^{-/-}$IL2Rg$^{-/-}$* recipients. (A)** The lists of DEGs between allografts from WT vs *Rag$^{-/-}$* **(Fig 4A, Fig S4A)** and WT vs *Rag$^{-/-}$IL2Rg$^{-/-}$* **(Fig 4B, Fig S4B)** were compared to identify shared and unique DEGs between the two comparisons. **(B-D)** Pathway analysis of the gene sets identified in **(A)** using the Enrichr[23-25] tool and Reactome[26] pathways database. The top ten most enriched pathways for each gene set were identified and plotted based on their adjusted p-values. **(B)** Enriched pathways from WT vs *Rag$^{-/-}$* unique DEGs. **(C)** Enriched pathways from shared DEGs. **(D)** Enriched pathways from WT vs *Rag$^{-/-}$IL2Rg$^{-/-}$* unique DEGs.

### SUPPLEMENTAL FIGURE 6

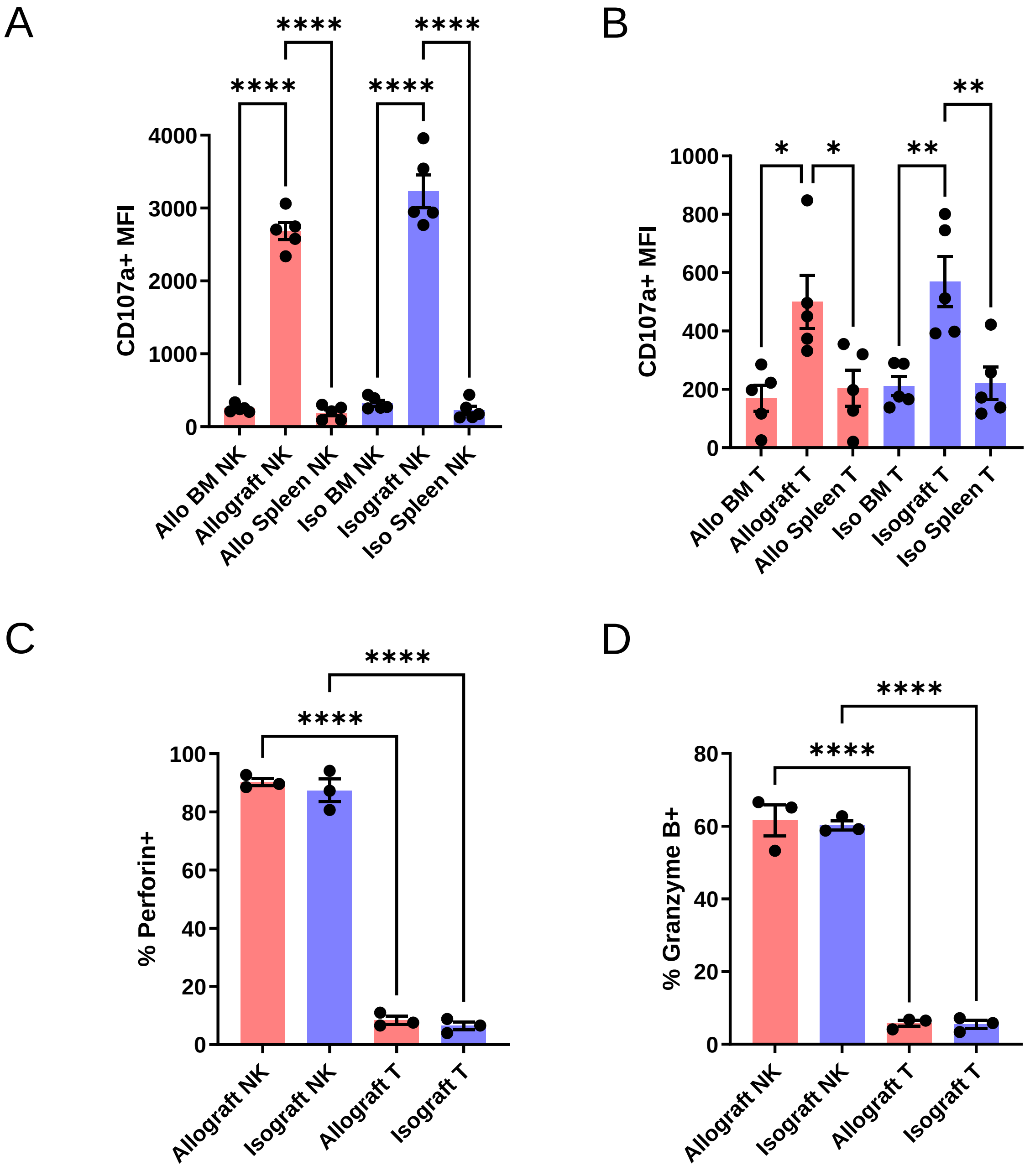

**Supplemental Figure 6. Additional analyses of recipient NK and T cells related to Fig 5.** Groups of wild type A/J and B6 hearts were transplanted to wild type B6 recipients. Grafts, recipient bone marrow and recipient spleens were harvested on day 1 post-transplant. **(A-B)** Summary data related to Fig 5D-E comparing the surface expression levels of CD107a on NK cells **(A)** and T cells **(B)** from grafts, recipient bone marrow, and recipient spleens. N=5 per group. **(C-D)** Proportion of perforin **(C)** and granzyme B **(D)** expressing allograft and isograft infiltrating NK and T cells from the experiment in **Fig 5F-G** (N=3 per group). Data reported as mean ± SEM, ✱ = $p<0.05$, ✱✱ = $p<0.01$, ✱✱✱ = $p<0.001$, ✱✱✱✱= $p<0.0001$

**Major Resources Table**

#### Animals (in vivo studies)

| Species | Vendor or Source | Background Strain | Sex | Persistent ID / URL |
|---|---|---|---|---|
| Mus musculus | The Jackson Laboratory | C57BL/6J | M | RRID: IMSR_JAX:000664 |
| Mus musculus | The Jackson Laboratory | A/J | M | RRID: IMSR_JAX:000646 |

#### Genetically Modified Animals

| | Species | Vendor or Source | Background Strain | Persistent ID / URL |
|---|---|---|---|---|
| B6.129S7-*Rag1*$^{tm1Mom}$/J (Rag1 KO) | Mus musculus | The Jackson Laboratory | C57BL/6J | RRID: IMSR_JAX:002216 |
| C57BL/6NTac.Cg-*Rag2*$^{tm1Fwa}$ *Il2rg*$^{tm1Wjl}$ (Rag2/Il2rg Double Knockout) | Mus musculus | Taconic Biosciences | C57BL/6NTac | RRID: IMSR_TAC:4111 |

#### Antibodies

| Target antigen (All anti-mouse) | Vendor or Source | Catalog # | Working concentration | Persistent ID / URL |
|---|---|---|---|---|
| CD45 RB705 | BD Biosciences | 570291 | 1:100 | RRID: AB_3086726 |
| CD90.2/Thy1.2 BV421 | Biolegend | 140327 | 1:100 | RRID: AB_2686992 |
| CD3 Spark 718 | Biolegend | 100282 | 1:100 | RRID: AB_2924440 |
| CD4 BV711 | BD Biosciences | 563726 | 1:100 | RRID: AB_2738389 |
| CD8 BV650 | BD Biosciences | 563234 | 1:100 | RRID: AB_2738084 |
| NK1.1 BV786 | Invitrogen | 417-5941-82 | 1:100 | RRID: AB_2925747 |
| CD335/NKp46 PE | Biolegend | 137604 | 1:50 | RRID: AB_2235755 |
| CD49b/DX5 APC | Biolegend | 108910 | 1:50 | RRID: AB_313417 |
| CD107a/LAMP-1 FITC | Biolegend | 121605 | 1:100 | RRID: AB_572006 |
| CD45R/B220 RY775 | BD Biosciences | 571753 | 1:100 | RRID: AB_3678612 |
| CD45 APC Fire750 | Biolegend | 103154 | 1:100 | RRID: AB_2572116 |
| CD11b BV421 | Biolegend | 101236 | 1:100 | RRID: AB_11203704 |
| Ly6G APC | Biolegend | 127614 | 1:100 | RRID: AB_2227348 |
| Ly6C R718 | BD Biosciences | 569482 | 1:100 | RRID: AB_3678613 |
| CD115/CSF-1R BV605 | Biolegend | 135517 | 1:100 | RRID: AB_2562760 |
| CD101 PECy7 | Invitrogen | 25-1011-82 | 1:100 | RRID: AB_2573378 |
| CD184/CXCR4 BV711 | Biolegend | 146517 | 1:100 | RRID: AB_2687244 |
| F4/80 RB705 | BD Biosciences | 570288 | 1:100 | RRID: AB_3678614 |
| dcTRAIL-R1/TNFRSF23 biotin | R&D Systems | BAM2378 | 1:75 | RRID: AB_2207073 |
| IFIT1 AF488 | Novus Biologicals | NBP2-71005AF488 | 1:50 | RRID: AB_3363105 |
| CD45 BV421 | BD Biosciences | 563890 | 1:100 | RRID: AB_2651151 |
| Granzyme B FITC | Biolegend | 515403 | 1:100 | RRID: AB_2114575 |
| Perforin AF647 | Biolegend | 154308 | 1:100 | RRID: AB_2922480 |
| IFNγ RY775 | BD Biosciences | 571773 | 1:100 | RRID: AB_3678615 |
| Ly6G APCCy7 | Biolegend | 127624 | 1:100 | RRID: AB_10640819 |
| CD45 APC | BD Biosciences | 559864 | 1:100 | RRID: AB_398672 |
| Mouse Fc Block (anti-CD16/32) | BD Biosciences | 553141 | 1:50 | RRID: AB_394656 |

DOI [to be added]

**Other**

| Description | Source / Repository | Persistent ID / URL |
|---|---|---|
| Streptavidin-PE (405204) | Biolegend | https://www.biolegend.com/en-us/products/pe-streptavidin-1475 |
| Ringer's solution (R5310-01) | B. Braun Medical Inc. | https://www.bbraunusa.com/en/products/b0/ringer-s-irrigationusp1000ml.html |
| 10X RBC Lysis Buffer (420301) | Biolegend | https://www.biolegend.com/en-us/products/rbc-lysis-buffer-10x-1498 |
| Collagenase D (11088882001) | Roche | https://www.sigmaaldrich.com/US/en/product/roche/colldro |
| DNase I (10104159001) | Roche | https://www.sigmaaldrich.com/US/en/product/roche/10104159001 |
| Brilliant Stain Buffer Plus (566385) | BD Biosciences | https://www.bdbiosciences.com/en-us/products/reagents/flow-cytometry-reagents/research-reagents/buffers-and-supporting-reagents-ruo/brilliant-stain-buffer-plus.566385 |
| HyClone Fetal Bovine Serum (SH30070.03) | Cytiva | https://www.cytivalifesciences.com/en/us/shop/cell-culture-and-fermentation/sera/fetal-bovine-serum/hyclone-defined-fetal-bovine-serum-fbs-u-s-origin-p-06090 |
| LIVE/DEAD Fixable Aqua (L34965) | ThermoFisher Scientific | https://www.thermofisher.com/order/catalog/product/L34965 |
| eBioscience Foxp3/Transcription Factor Staining Buffer Set (00-5523-00) | ThermoFisher Scientific | https://www.thermofisher.com/order/catalog/product/00-5523-00 |
| eBioscience Cell Stimulation Cocktail plus protein transport inhibitors (00-4975-93) | ThermoFisher Scientific | https://www.thermofisher.com/order/catalog/product/00-4975-93 |
| RNeasy Fibrous Tissue Mini Kit (74704) | QIAGEN | https://www.qiagen.com/us/products/discovery-and-translational-research/dna-rna-purification/rna-purification/total-rna/rneasy-fibrous-tissue-mini-kit |
| nCounter Mouse PanCancer Immune Profiling Panel | NanoString, Bruker Spatial Biology | https://nanostring.com/products/ncounter-assays-panels/oncology/pancancer-immune-profiling/ |
